# Mapping differences in mammalian distributions and diversity using environmental DNA from rivers

**DOI:** 10.1101/2021.04.27.441610

**Authors:** Holly A. Broadhurst, Luke M. Gregory, Emma K. Bleakley, Joseph C. Perkins, Jenna V. Lavin, Polly Bolton, Samuel S. Browett, Claire V. Howe, Natalie Singleton, Darren Tansley, Naiara Guimarães Sales, Allan D. McDevitt

## Abstract

**Aim:** Finding more efficient ways to monitor, and estimate the diversity of, mammalian communities is a major step towards their management and conservation. Environmental DNA (eDNA) from river water has recently been shown to be a viable method for biomonitoring mammalian communities. Yet, most of the studies to date have focused on the potential for eDNA to detect individual species, with little focus on describing patterns of community diversity and structure. In this study, we focus on the sampling effort required to reliably map the diversity and distribution of semi-aquatic and terrestrial mammals and allow inferences of community structure surrounding rivers.

**Location:** Southeastern England

**Methods:** We used eDNA metabarcoding on water samples collected along two rivers and a beaver enclosure over two days, targeting terrestrial and semi-aquatic mammals. Mammalian community diversity and composition was assessed based on species richness and β-diversity. Differences between river communities were calculated and partitioned into nestedness and turnover, and the sampling effort required to rapidly detect semi-aquatic and terrestrial species was evaluated based on species accumulation curves and occupancy modelling.

**Results:** eDNA metabarcoding efficiently detected 25 wild mammal species from five orders in two days of sampling, representing the vast majority (82%) of the species expected in the area. The required sampling effort varied between orders, with common species (generally rodents, deer and lagomorph species) more readily detected, with carnivores detected less frequently. Measures of species richness differed between rivers (both overall and within each mammalian order) and patterns of β-diversity revealed the importance of species replacement in sites within each river, against a pattern of species loss between the two rivers.

**Main conclusions:** eDNA metabarcoding demonstrated its capability to rapidly detect mammal species, allowing inferences of community composition that will better inform future sampling strategies for this Class. Importantly, this study highlights the potential use of eDNA data for investigating mammalian community dynamics over different spatial scales.

## 1. Introduction

Mammalian populations have suffered significant declines globally, with one in four species believed to be threatened (defined as critically endangered, endangered or vulnerable; IUCN, 2021). Information on species’ distributions is therefore critical to support effective evidence-based management (Mathews et al., 2018). However, undertaking surveys to capture a broad range of mammals within a particular area or region can be logistically challenging in terms of effort, cost and time (Garden et al., 2007). This is especially evident for species that are difficult to visually encounter or which occur at low densities. Mammals are traditionally surveyed using a variety of methods including camera trapping, live-trapping and/or field sign surveys (Sales, McKenzie et al., 2020) with the accuracy of these methods heavily reliant on the intensity of sampling efforts and the susceptibility of species and individuals to capture/detection by each method. Each method may have additional concerns or limitations, such as ethical considerations in live-trapping (Sikes et al., 2016), surveyor expertise in correctly identifying field signs (Harrington et al., 2010) and camera trap placement (Littlewood et al., 2021; Kaizer et al., 2021). Given the wide variety of ecologies exhibited within mammals, there is clearly no ‘one size fits all’ method for monitoring either the entire or a significant component of the overall mammalian community.

The emergence of environmental DNA (eDNA) as both a viable and reliable method for biomonitoring is rapidly transforming how species and community-wide surveys are undertaken (Deiner et al., 2017; Fediajevaite et al., 2021). eDNA is any genetic material that has been shed into the environment by macro-organisms through sloughed skin cells, blood, faeces/urine and saliva with no obvious signs of biological source material (Pawlowski, Apotheloz-Perret-Gentil & Altermatt, 2020). When eDNA is combined with next-generation sequencing (NGS) technology via DNA metabarcoding with universal primers, it has the potential to facilitate rapid biodiversity assessments in diverse and complex ecosystems as it can identify multiple species simultaneously from one environmental sample (Deiner et al., 2017). Since water has been shown to be a reliable source of eDNA (Deiner et al., 2017), most eDNA metabarcoding applications to date on vertebrates have been focused on monitoring fishes and amphibians (e.g. McDevitt et al., 2019; Valentini et al., 2016). However, recent studies have demonstrated that eDNA retrieved from water from both lotic (Sales, McKenzie et al., 2020; Sales, Kaizer et al., 2020; Lozano & Caballero, 2021; Macher et al., 2021; Mariani et al., 2021) and lentic (Ushio et al., 2017; Harper et al., 2019) systems can detect components of the overall terrestrial and semi-aquatic mammalian community.

Mammals frequently come into contact with water through behaviours such as drinking, bathing, foraging, urinating and defecating either in or near the water system (Rodgers & Mock, 2015; Williams et al., 2018) and even mammalian species that display limited interactions with water have been shown to be effectively detected by eDNA (Williams et al., 2018; Sales, Kaizer et al., 2020). Comparisons between eDNA and camera trapping surveys have revealed considerable overlap between the methods in terms of the species detected (Sales, McKenzie et al., 2020; Sales, Kaizer et al., 2020; Harper et al., 2019; Leempoel et al., 2020) or found comparable detection probabilities from eDNA and field surveys (Sales, McKenzie et al., 2020; Lugg et al., 2018). Most of the studies to date have focused on the potential for eDNA to detect individual mammalian species (Ushio et al., 2017; Sales, McKenzie et al., 2020), with little to no focus on describing patterns of community diversity and structure. This is perhaps unsurprising given the infancy of eDNA in terms of its application for monitoring/surveying mammals (Sales, McKenzie et al., 2020; Sales, Kaizer et al., 2020).

Biodiversity estimates obtained from eDNA metabarcoding are known to be influenced by sampling effort, with some taxonomic groups requiring greater spatiotemporal coverage of eDNA sampling (e.g. carnivores; Harper et al., 2019; Leempoel et al., 2020; Sales, McKenzie et al., 2020). Quantifying the differences between mammalian communities is a major step in understanding the factors that shape communities. The number of species in a local assemblage can help indicate the health of a local ecosystem, with the presence of certain species, such as otters (*Lutra lutra*; Esposito et al., 2020), providing critical insights into factors that affect environmental health (e.g., pollution). Riverine ecosystems are among the most dynamic habitats which support a rich diversity of species but are also exposed to multiple threats including pollution, the spread of invasive species, habitat fragmentation and degradation. Due to the connectivity of these freshwater systems, these threats are easily transported and have profound effects on the distribution of biodiversity (Collen et al., 2009; Dudgeon et al., 2019). As a result of rivers’ roles as ‘conveyor belts of biodiversity information’ (Deiner et al., 2016), they represent suitable sampling points for inferring the distribution of mammalian communities with highly divergent functional adaptations (Sales, McKenzie et al., 2020).

To understand processes responsible for shaping community assembly, estimating species richness and β-diversity (also referred to as inter-community structure) is of paramount importance. β-diversity can be partitioned and used to determine if communities are subsets of sites with higher species richness (nestedness), or if the dissimilarity between sites is driven by species replacement (i.e., spatial turnover; Baselga, 2010). In this context, an eDNA-based ecological assessment can contribute to the identification of locations that require protection and direct future conservation management (Socolar et al., 2016). The main aim of this study is to explore the use of eDNA metabarcoding for assessing the distribution and diversity of terrestrial and semi-aquatic mammals around two adjacent rivers. eDNA sampling was conducted in transects along the Rivers Colne and Blackwater in Essex (Fig. 1A) where the Essex Wildlife Trust (EWT) have implemented species management and habitat improvements, including both re-introduction and eradication programmes for critically endangered and invasive mammals, respectively. Additionally, eDNA sampling was conducted in and downstream of a nearby and newly established beaver (*Castor fiber*) enclosure (Fig. 1B) to determine its efficiency for detecting the focal species and surrounding mammalian community. Aiming to expand upon the application of eDNA for monitoring mammalian communities, our objectives are to determine the optimal sampling effort to (1) adequately describe overall mammalian species diversity, and within each mammalian order identified, from eDNA recovered from river water, and to (2) quantify the differences among mammalian communities through analysing spatial patterns of β-diversity within and between rivers.

**Figure 1.**
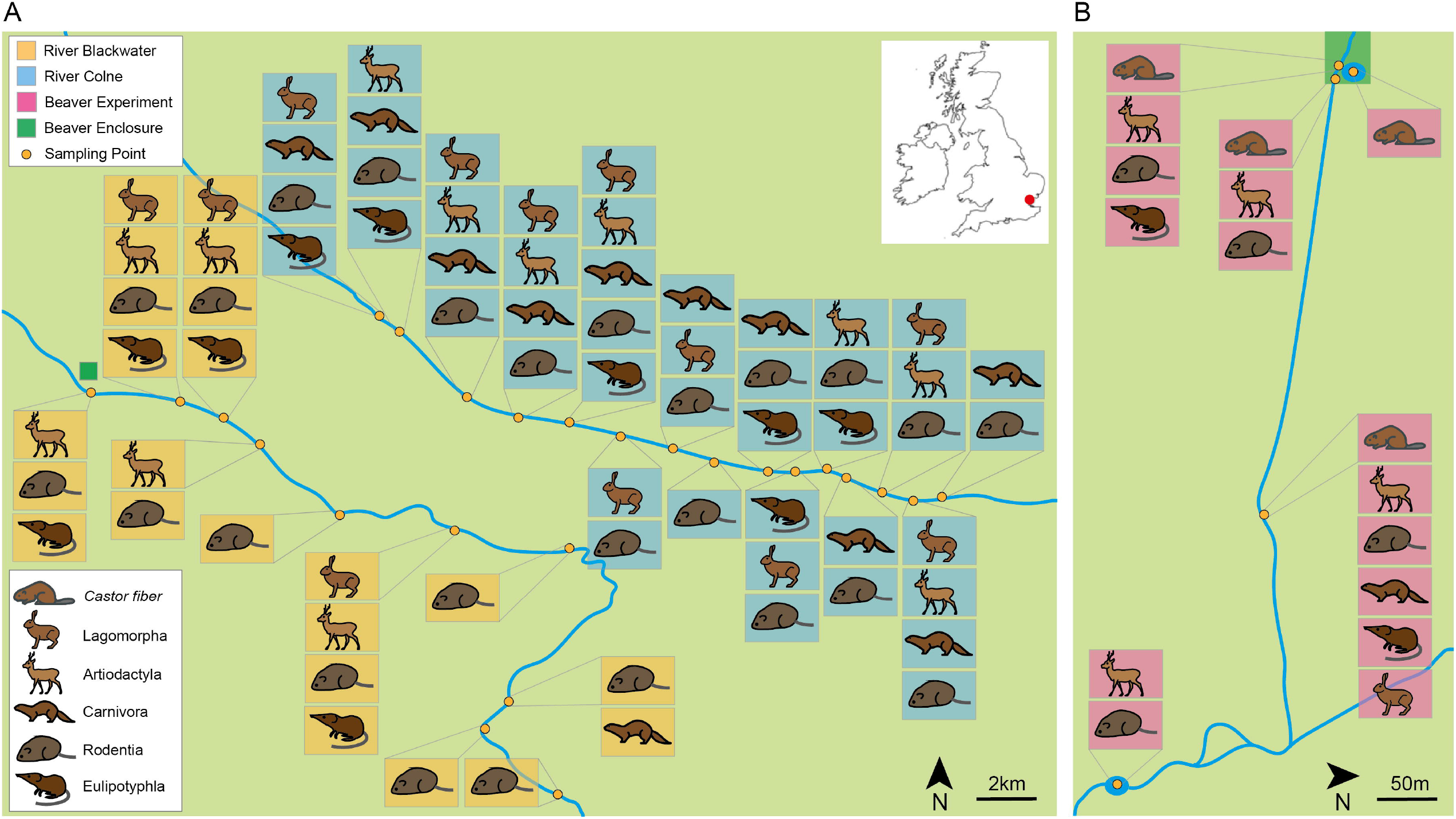
Community structure of each sampling site represented by Order for the Rivers Blackwater (orange) and Colne (blue; A) in Essex, England (approximate location shown in red on the inset map). For the beaver experiment (pink; B), beaver (*Castor fiber*) detections specifically are also shown.

## 2. Methods

### 2.1 Field sampling

Samples were collected in Essex, England along the Rivers Colne and Blackwater (Figs. 1A and S1A). Five 500 ml water replicates were collected from 15 sites along the River Colne and 10 sites along the River Blackwater on 29th–30th July 2019 at roughly equal intervals within each river and accounting for access. In this area, the EWT implemented the Essex Water Vole Recovery Project (EWVRP) in 2007, focusing on reintroducing water voles (*Arvicola amphibius*) to rivers in Essex, targeted intensive control of the invasive American mink (*Neovison vison*) and habitat improvements (McGuire & Whitfield, 2018). Mink removal has resulted in the natural recolonization of water voles across over 500km^2^ of North East Essex and the River Colne Water Vole Translocation Project (RCWVTP) was implemented in 2009, with 600 water voles released on the Colne between 2010-2012. Mink control has occurred on the river Blackwater but no water vole releases have taken place. In addition to the aforementioned sites, eDNA sampling was conducted at Spains Hall Estate (Figs 1B and S1B), where a pair of Eurasian beavers (*C. fiber*) are housed in a fenced enclosure outdoors. The beavers were released into the enclosure in March 2019. For the ‘beaver experiment’, sampling deviated slightly from the two main rivers described above in that four water replicates were collected from each of the three sampled locations in the beaver enclosure (B1-B3; where B1 was a pond within the enclosure and B2-3 along the stream adjacent to it), five replicates were collected from downstream of the beaver enclosure (B4) and eight replicates were collected from a large pond inlet of a brook (B5; Fig. S1B).

Water samples were collected using sterile 500 ml water bottles on the shoreline at a reachable distance with complete submersion beneath the surface (Fig. S2). Four field controls consisting of a bottle of distilled water (500 ml) were opened briefly at the beginning and end of each of the two sampling days to test for cross-contamination during sampling. Samples and field blanks were transported in cool boxes, sterilised with 10% bleach and 70% ethanol. Samples were filtered on the same day as collection using sterilised single-use syringes and 0.45 μm Sterivex filter (Merck Millipore, Darmstadt, Germany). The filters were stored at 4°C for 2 days prior to transportation to the laboratory, transported in cool boxes with ice packs and then stored at -20°C until DNA extraction. Water sampling and filtration followed the same procedures as described above.

### 2.2 eDNA extraction and metabarcoding

DNA was extracted from the filters in a dedicated eDNA clean room following the Mu-DNA protocol (Sellers et al., 2018). Field controls were extracted first, followed by the eDNA samples. Five DNA extraction negative controls (one for each day of extractions) containing only extraction buffers were also included. All surfaces were sterilised with 10% bleach and then washed with 70% ethanol. Small tools were placed in a UV Stratalinker® before, in-between and after extracting each sample to reduce the risk of cross-contamination.

DNA extracts were stored at -20°C until PCR amplification. Eluted eDNA was amplified using the MiMammal 12S primer set (MiMammal-U-F, 5′-GGGTTGGTAAATTTCGTGCCAGC-3′; MiMammal-U-R, 5′-CATAGTGGGGTATCTAATCCCAGTTTG-3′; Ushio et al., 2017) targeting a ∼170bp amplicon from a variable region of the 12S rRNA mitochondrial gene with sample-specific multiplex identifier (MIDs) tags. PCR amplification protocols followed Sales, McKenzie et al. (2020). PCRs were conducted in triplicates to reduce bias in individual reactions and the replicates were pooled prior to library preparation. Amplification was validated using 1.2% agarose gel electrophoresis stained with GelRed (Cambridge Bioscience). In total, 185 samples were analysed, including 158 eDNA samples, four field collection blanks, five extraction blanks, 10 PCR negative controls and eight PCR positive controls (i.e. DNA extraction from a non-target species that is not locally present, the northern muriqui *Brachyteles hypoxanthus* from Brazil, at a concentration of 0.05 ng/µL). eDNA samples were equally distributed into two sequencing libraries, with replicated extraction controls in each library. A left-sided size selection was performed using 1.1x Agencourt AMPure XP (Beckman Coulter) and Dual-Index adapters (Illumina) were added to each library using KAPA HyperPrep kit (Roche). Each library was then quantified by qPCR using NEBNext qPCR quantification kit (New England Biolabs) and pooled in equimolar concentrations. Libraries were sequenced using an Illumina MiSeq v2 Reagent Kit for 2×150 bp paired-end reads (Illumina, San Diego, CA, USA).

### 2.3 Bioinformatic analysis

The bioinformatic analysis was conducted using OBITools metabarcoding package (Boyer et al., 2016) following the protocol described in in Sales, McKenzie et al. (2020). Briefly, the quality of the reads were assessed using FASTQC (Andrews, 2015), a filter was used to select fragments of 140-190bp and to remove reads with ambiguous bases using obigrep, followed by a sequence clustering using SWARM (Mahé et al., 2015) and a taxonomic assignment conducted using ecotag against a custom database (Sales, McKenzie et al., 2020). An additional conservative filtering procedure was conducted to exclude MOTUs/reads originating from putative sequencing errors or contamination in order to avoid false positives (Table S1). First, to account for the occurrence of tag-jumps between tagged amplicons (Schnell et al., 2015), the frequency of tag-jumping was calculated for each sequencing library by dividing the total number of reads of the positive control (PC, *B. hypoxanthus*) recorded in the actual eDNA samples and negative controls by the total number of reads of the PC in the PC samples. The frequency was taken off all MOTUs and the PC was removed. Then, to remove putative contaminants, the maximum number of reads recorded for a MOTU in one of the negative controls (whether this be a field collection blanks, extraction blanks or PCR negative controls) was removed from all samples for each MOTU. Finally, non-target MOTUs (non-mammal species, human and domestic species) and MOTUs that were likely to have been carried over from contamination were discarded from the dataset by removing MOTU’s with <5 total reads, and only MOTUs that were identified at species level with a best identity of > 0.98 were included (Sales, McKenzie et al., 2020).

### 2.4 Statistical analysis

All statistical analyses were performed using R v4.0.0 (R Core Team, 2020). After bioinformatic filtering, bubble charts were created using *ggplot2* (Wickham & Chang, 2016) showing the proportional read count of each species identified at each sampling site along each river and around the beaver experiment. Read counts in a water replicate at a sampling site were then converted into binary presence-absence data for downstream analyses. As we aimed to compare the mammalian diversity present in the two river systems (Colne and Blackwater), for subsequent analyses, a dataset comprising only the 15 sites from the River Colne and 10 sites from the River Blackwater (i.e., excluding sites B1-B5 representing the beaver experiment as these included ponds) was used. Species accumulation curves were created, using the R package *iNext* (Hsieh & Chao, 2020), to determine if the number of sample sites sampled was adequate to represent the overall species diversity along both rivers and when observing the systems as a whole, and to estimate the sample effort needed to fully determine the species richness (Hsieh, Ma, & Chao, 2016). Mathews et al. (2018) was used to infer known mammalian species (excluding bats) distributions in the region from 1995-2016.

In order to determine detection probabilities of each species’ eDNA, a single season occupancy model (MacKenzie et al., 2002) was applied to the data where detection histories were created using each of the five water replicates taken at a site as sampling occasions (MacKenzie et al., 2017), following Sales, McKenzie et al. (2020). The core assumption here is that the underlying occupancy state (i.e. occupied or empty) is constant over the sampling period, and therefore, every sampling occasion is an imperfect observation of the true occupancy status of a species’ eDNA at that site. Our primary aim was to compare eDNA detectability across different species within our sampling effort, so we did not consider any other competing models (Sales, McKenzie et al., 2020). These analyses were conducted separately for each species, overall and within each river (excluding when a species’ eDNA was not detected in a particular river), using the R package *unmarked* (Fiske & Chandler, 2011).

To illustrate the differences in the average species richness in sample sites between the two river systems, box and jitter plots were created using the *tidyverse* R package (Wickham et al., 2019). A non-metric multidimensional scaling (NMDS) analysis was completed to visualise the differences in community composition between both rivers using the metaMDS function from the *vegan* R package (Oksanen et al., 2019). Jaccard index distances were utilised to create the NMDS plot with a stress value being calculated to verify the goodness-of-fit between the NMDS ordinations and a commonly accepted set of guidelines (Dexter, Rollwagen-Bollens & Bollens, 2018). Differences between the two rivers were calculated using a permutational multivariate analysis of variance (PERMANOVA) with 1000 permutations being performed using the adonis function in the *vegan* R package (Oksanen et al., 2019) applying Jaccard index distances.

The *betapart* package (Baselga et al., 2012) was used to assess the spatial patterns of β-diversity using multiple-site dissimilarity measures across all analysed sites. β-diversity was calculated and partitioned into nestedness (i.e., species loss) and turnover (i.e., species replacement) components using the Sørensen index, following Baselga (2010). Three multiple-site dissimilarities were estimated including βsor (Sørensen dissimilarity), βsim (Simpson dissimilarity, turnover component of Sørensen dissimilarity) and βsne (nestedness component of Sørensen dissimilarity) using three different datasets (one for each river -Blackwater and Colne, and one combining the data for both rivers).

## 3. Results

### 3.1. Species detections using eDNA metabarcoding

The MiSeq sequencing run yielded a total of 12,021,106 raw reads. Following filtering criteria, 2,447,689 reads were retained (Table S1). In total, 25 wild mammal species were detected across all sampling locations (Table S2), 23 species on the River Colne (Figs 2 and S3), 12 species on the River Blackwater (Figs 2 and S4) and 12 species in and downstream of the beaver enclosure (Fig. S5). Overall, mammals were detected from five orders: Artiodactyla (3 species), Carnivora (7 species), Eulipotyphla (4 species), Lagomorpha (2 species) and Rodentia (9 species). This comprised ten families and twenty genera (Figs 1 and 2; Table S2). This list included 15 species designated as Least Concern, two Endangered, one Critically Endangered, three naturalised and four non-native (Table S2; Crawley et al., 2020). The Chao II estimation based on the eDNA results predicted 27 species (95% CI: 24-41; Table S3) overall and this is a close representation of the 28 terrestrial and semi-aquatic mammal species expected in Essex (Mathews et al., 2018). On the River Colne 23 species were detected, with 28 predicted (95% CI: 24-49; Table S3) according to the Chao II estimate, whereas 12 species were predicted (95% CI: 12-20; Table S3) on the river Blackwater which represents the same number of species observed. Of the 25 species identified overall with eDNA, 23 are known from the region from 1995-2016 records (Mathews et al., 2018), with the beaver being reintroduced after this period. One species, the red squirrel (*Sciurus vulgaris*), was last detected in the region in 1971 but the species was reintroduced to Mersea Island in 2012 (Dobson & Tansley, 2014), approximately 10km from where it was detected using eDNA. For completeness, this detection was retained in the downstream analyses.

**Figure 2.**
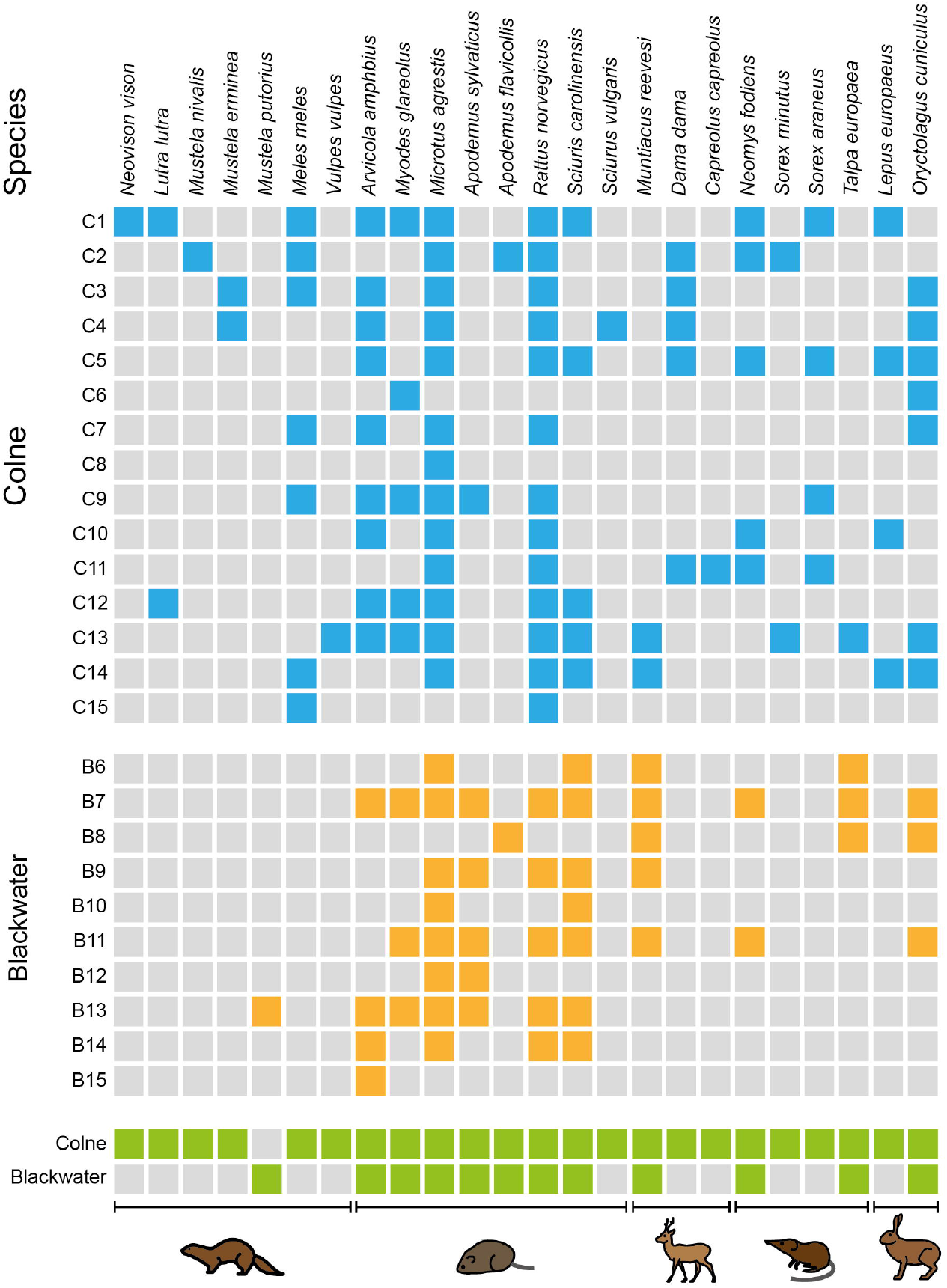
Species detections from eDNA metabarcoding data by sampling site (see Fig. S1A) within the Rivers Colne (blue) and Blackwater (orange), and combined (green). Species are grouped by order (left to right: Carnivora, Rodentia, Artiodactyla, Eulipotyphla and Lagomorpha).

Species from the order Rodentia were the most prevalent, with at least one species from this order being detected at all sampling sites (Fig. 1). Ten unique species were detected on the river Colne including five carnivores: European otter (*Lutra lutra*), European badger (*Meles meles*), stoat (*Mustela erminea*), least weasel (*Mustela nivalis*) and American mink (*N. vison*); two species from the order Artiodactyla: fallow deer (*Dama dama*) and roe deer (*Capreolus capreolus*); common shrew (*Sorex araneus*) from the order Eulipotyphla; red squirrel (*S. vulgaris*) from Rodentia and brown hare (*Lepus europaeus*) from Lagomorpha (Table S2). One unique carnivore was detected at one site on the river Blackwater (B13): European polecat (*Mustela putorius;* Figs 2 and S4). Eurasian beaver (*C. fiber*) was detected at all sampling sites inside the beaver enclosure (B1-B3) and approximately 300m downstream of the enclosure (B4) but was not detected at the large pond inlet (B5; Figs 1B and S1B; Fig. S5).

### 3.2. Occupancy and detection probabilities in the Rivers Colne and Blackwater

Based on the five water replicates taken at each sampling site, site occupancy and detection probabilities based on eDNA varied markedly between species. Six species (three from the order Carnivora, two from Rodentia and one from Artiodactyla) had detection probabilities close to zero given that they were detected in only a single water replicate at either one or two sampling sites (Figs 2 and 3; Table S4). Despite being detected at eight sampling sites (Fig. 2), the bank vole (*Myodes glareolus*) had a low detection probability of only 0.07 because it was generally only detected in a single water replicate at each site it was found. For the Carnivora, American mink (*N. vison*) and stoat (*M. erminea*) had high detection probabilities despite only being found at 1-2 sampling sites along the River Colne. This is due to both being detected in 2-4 replicates when found at a site. The badger (*M. meles*) was the most frequently detected carnivore (seven sites in the River Colne; Fig. 2), but had a similar detection probability (0.24 compared to 0.20) to the otter (*L. lutra*; Fig. 3; Table S4), which was detected at only two sites on the Colne.

**Figure 3.**
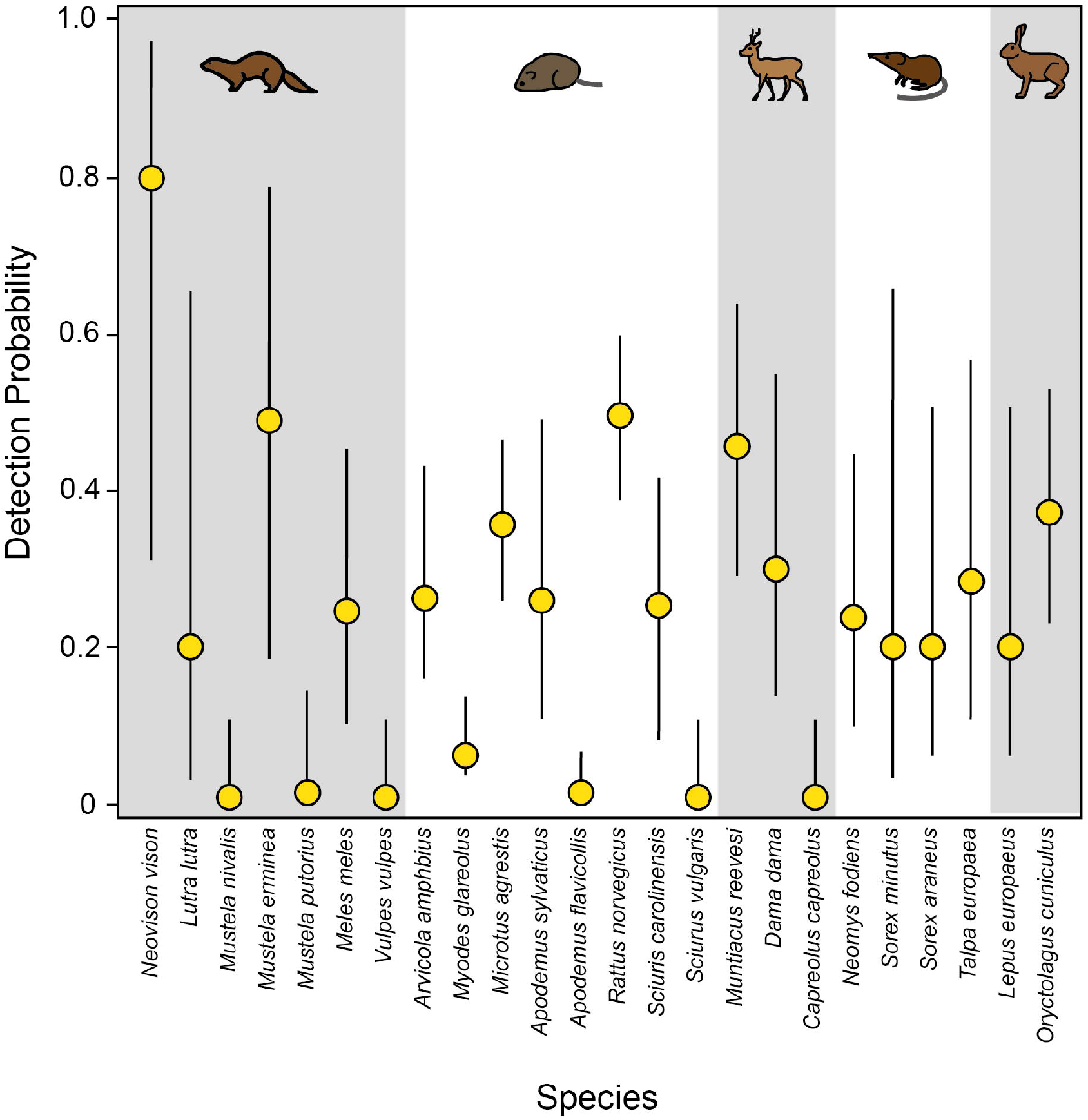
Estimated detection probabilities (yellow) for each species (grouped by order), with vertical lines representing the 95% confidence intervals.

Species from Rodentia were generally the most frequently detected species at sampling sites overall (e.g. field vole *Microtus agrestis* at 21/25 sites; brown rat *Rattus norvegicus* at 18/25 sites; water vole *A. amphibius* at 13/25 sites; grey squirrel *S. carolinensis* at 12/25 sites; *M. glareolus* at 8/25 sites and woodmouse *Apodemus sylvaticus* at 6/25 sites; Fig. 2; Table S2). These species were frequently detected in the beaver experiment also (Table S2; Fig. S5). Detection probabilities in the Colne and Blackwater combined ranged from 0.26 to 0.49 (with the exception of the aforementioned *M. glareolus*) for these frequently occurring species (Fig. 3; Table S4). From the order Artiodactyla, *C. capreolus* was only detected on a single occasion but *Muntiacus reevesi* and *D. dama* were detected at seven and five sampling sites, respectively. Detection probabilities were 0.46 and 0.30 for *M. reevesi* and *D. dama*, respectively. Of the species from Eulipotyphla, the water shrew (*Neomys fodiens*) was found at 7/25 sites, common shrew (*S. araneus*) and mole (*Talpa europaea*) at four sites and pygmy shrew (*S. minutus*) at two. Detection probabilities were similar across these species, ranging from 0.20 to 0.28. For Lagomorpha, the brown hare (*L. europaeus*) was found at four sites on the Colne (detection probability of 0.20), with the European rabbit (*Oryctolagus cuniculus*) detected at 10/25 sites and a detection probability of 0.37 (Fig. 3; Table S4). With the notable exceptions of *M. agrestis* and *R. norvegicus*, 95% confidence intervals were generally large however around these estimates for detection probabilities for most species (Fig. 3; Table S4).

### 3.3. Sampling site effort

Evaluation of the species accumulation curves and their asymptotes provide insights into the sampling effort needed to achieve a comprehensive view of the mammalian diversity around the sampling area. By visually examining the asymptote of the accumulation curve for the overall diversity of combined sites for the River Colne and Blackwater, the total of 25 sites herein achieved to capture 24 species (85%) of the 28 expected semi-aquatic and terrestrial mammals present in the area, and to reach the asymptote a total of 45 sampling sites may be required (Fig. 4). When analysing the rivers individually an increased sampling effort is required for the River Colne as the curve is gradually increasing toward the asymptote, the 15 sites detected 23 species (82%) out of the predicted 28 species that are known to inhabit the sampling area. The River Blackwater required less sampling effort due to the lower overall diversity of the river (Fig. 4).

**Figure 4.**
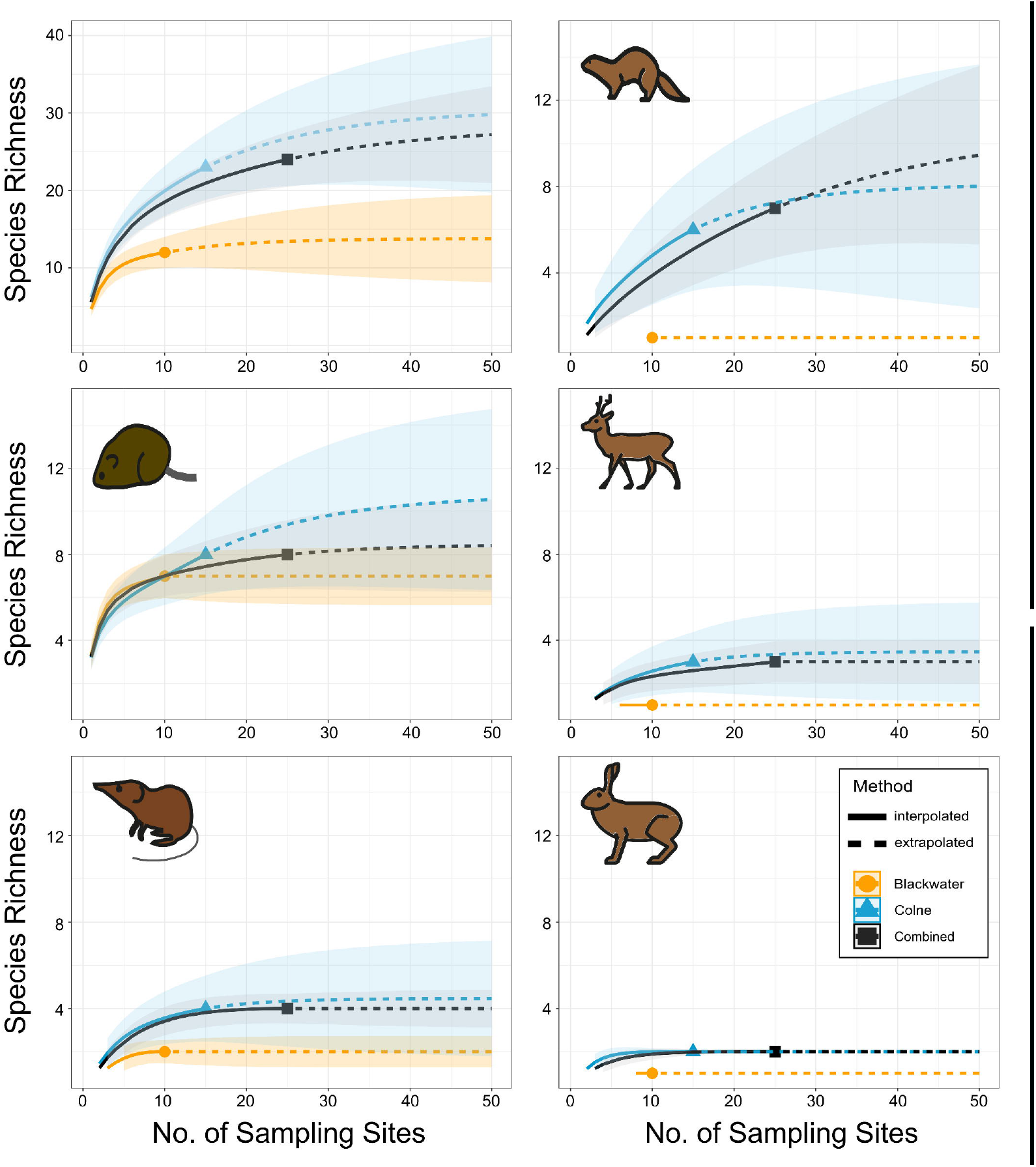
Accumulation curves of species detected according to the number of sampled sites, including data comprising all species together and divided by Order (represented by the symbols depicted in Fig. 1A) for each river (Blackwater in orange and Colne in blue) and both rivers combined (black). The number of species expected according to Mathews et al. (2018) is indicated by the red dotted line. From left to right and top to bottom: Overall, Carnivora, Rodentia, Artiodactyla, Eulipotyphla and Lagomorpha.

When visually inspecting the accumulation curves for each mammalian order for the river Blackwater, they indicate that sufficient sampling effort has been acquired for all orders as all the curves have reached an asymptote within our sampled sites. This is in contrast to the species accumulation curves on the river Colne where only the order Lagomorpha, representing two species, has plateaued within our sampled sites. The orders Eulipotypyla and Artiodactyla did not reach a plateau within our sampled sites but only one species of each order was not detected that are known in the sampling area (Fig. 4; Table S2; Mathews et al., 2018). The accumulation curves for Rodentia did not reach an asymptote and three species of the order known in the sampling area were not detected. Although all carnivores that are known in the sampling area have been detected using eDNA metabarcoding, the accumulation curve represents an over-prediction of species richness for this order. When the sample sites are combined for both rivers, the orders Artiodactyla, Eulipotyphla and Lagomorpha are shown to reach an asymptote within our sampled sites (Fig. 4).

### 3.4. Species richness and β-diversity

The River Colne had a higher species richness when compared to the Blackwater, both overall (Fig. 5A) and when analysing the species richness per taxonomic order (Fig. 4). Differences in community compositions between both river systems were initially visualised with NMDS ordination plots which demonstrates the sampling sites for each river grouped in two clusters, with a small overlap (Fig. 5B). Furthermore, this dissimilarity pattern was further corroborated with the PERMANOVA test which determined that communities were significantly different among the rivers, despite the low variance explained (R^2^ = 0.106, *p* = 0.002).

**Figure 5.**
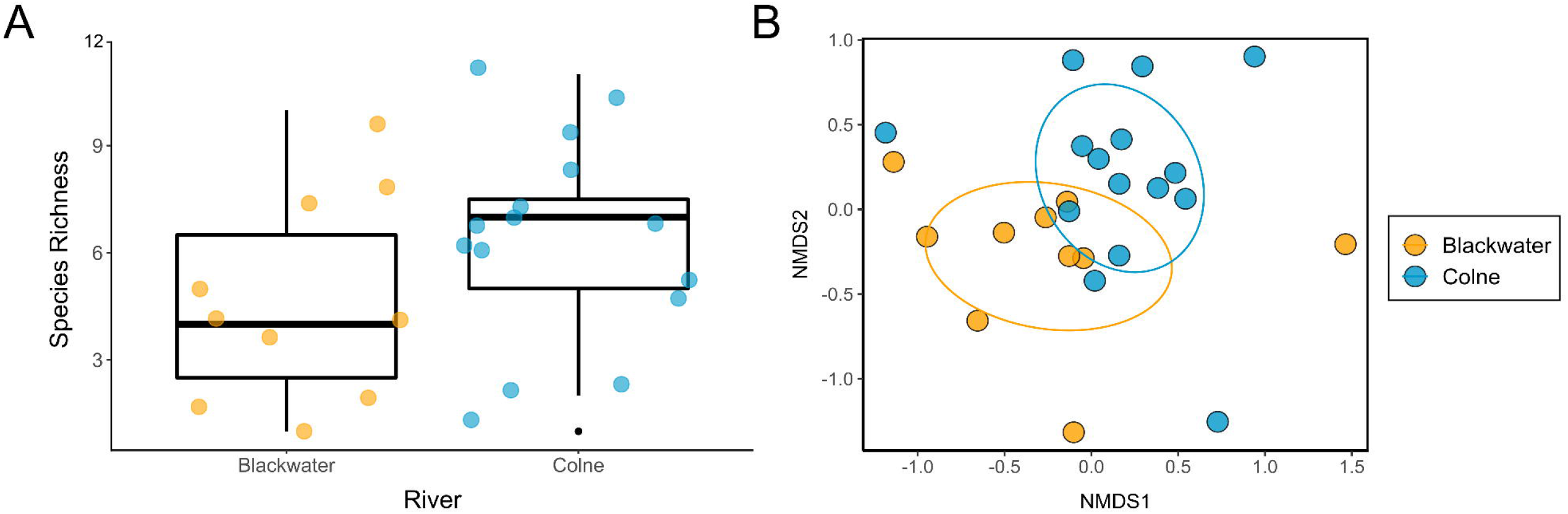
Box plot of overall species richness for each river (A) and an NMDS plot representing β-diversity based on Jaccard distances (B) of sampling sites on the Rivers Colne (blue) and Blackwater (orange).

Dissimilarity measures of the estimated overall community composition revealed high β-diversity for each river (Colne: βsor = 0.8376; Blackwater: βsor = 0.7690), with a lower dissimilarity when the rivers are compared (βsor = 0.3714). The compositional dissimilarity found in the River Colne was mainly associated with a high rate of species turnover with nestedness contributing a marginal amount (βsim = 0.7276, βsne = 0.1100), demonstrating that the β-diversity patterns are mostly caused by species replacement between sites. A similar pattern was found for the River Blackwater, but including an increase in the contribution of the nestedness component (βsim = 0.5395, βsne = 0.2295), with species replacement and species loss between sites both contributing to the high β-diversity values. In contrast to that, when comparing the assemblages of both rivers, most of the dissimilarity was due to nestedness (βsim = 0.0833, βsne = 0.2881).

## 4. Discussion

Due to the high costs in terms of effort and economics for monitoring entire communities, biodiversity assessments are often confined to very few indicator species and can therefore only provide a reduced representation of overall community dynamics and ecosystem health (Hilty & Merenlender, 2000). eDNA metabarcoding enables large-scale and multi-taxa surveys from material that can be collected rapidly in the field, and this multi-species monitoring could lead to more effective ecosystem-wide biodiversity assessments (Deiner et al., 2017). The mainstream application of eDNA for monitoring mammalian communities is certainly recent and have so far been relatively limited in scale (e.g. Harper et al., 2019; Sales, McKenzie et al., 2020; Sales, Kaizer et al., 2020). Sales, McKenzie et al. (2020) highlighted the potential of eDNA metabarcoding in riverine ecosystems to provide a rapid and rough ‘distribution map’ for mammalian biodiversity at a landscape scale. Here we expand upon this and previous studies by incorporating more intensive sampling in two major and adjacent rivers in southeastern England to investigate the effort required to capture the mammalian community, elucidate patterns of diversity and quantify differences within and among these sampled waterways.

In terms of providing a reliable snapshot of the mammalian community, this current study has demonstrated the power of eDNA-based monitoring for ‘capturing’ almost the entire known terrestrial and semi-aquatic mammalian community in the area in ∼30 hours of field sampling. 23 of the 28 known mammals in the area were detected in both rivers (Mathews et al., 2018). Considering the species accumulation curves, most of the mammalian diversity present in the surroundings of both rivers has been detected by eDNA (Fig. 4). Despite the potential need for an increased sampling effort to detect all species in the area, only five species expected in the region were not identified in this study (Table S2). The sampling period here represents a much shorter collection period than would be needed to fully estimate species composition by camera trapping, field signs and physical sightings (Roberts, 2011; Abrams et al., 2019). With 82% of the expected species detected here using eDNA (plus two recent additions), this is on the upper end of what camera trapping is expected to capture (57-86% of terrestrial mammals and avian species; Boitani, 2016). It is clear however that considerable effort is still required in terms of the number of sites sampled and replicates taken within a small geographic area for eDNA-based monitoring of mammals (Figs 1 and 2; Macher et al., 2021). A total of eight species were detected in only one or two locations (Fig. 2; Table S2) and most species had estimated detection probabilities of ∼0.20 (Fig. 3, Table S4), suggesting that all five water replicates are generally necessary to detect even common species. An alternative approach would be to filter larger volumes of water within a river system (Cantera et al., 2019; Bessey et al., 2020) but this would limit the applications of this method over larger spatial scales (because of the specialized pumps/equipment required; Cantera et al., 2019) and the involvement of citizen scientists to undertake the sampling (something that is now being applied more widely in eDNA-based monitoring; Biggs et al., 2015). The main consideration around the approach used here is one that can be adopted for national-based monitoring schemes with the involvement of citizen scientists and local conservation groups and this study has clearly shown that such an approach is logistically feasible.

A clear advantage of eDNA-based surveys in contrast to other sampling techniques (e.g. camera traps) is that it does not appear to be affected by animal body mass, since it has been shown to be capable of capturing effectively and simultaneously small to large mammals. With few exceptions, most species within the orders Artiodactyla, Eulipotyphla, Lagomorpha and Rodentia were frequently detected at multiple sites each (Figs 1 and 2) and had more consistent detection probabilities across species (Fig. 3; Table S4). Five species from Rodentia (*A. amphibius, M. glareolus, M. agrestis, R. norvegicus* and *S. carolinensis*), and one each from Artiodactyla (*M. reevesi*), Eulipotyphla (*N. fodiens*), Lagomorpha (*O. cuniculus*) were detected at multiple sites in each river system (Fig. 2). This generally reflects species which are known to be abundant in the region and/or group-living, factors which are important for eDNA detections (Sales, McKenzie et al., 2020; Williams et al., 2018). Other species such as the elusive water shrew (*N. fodiens*) is semi-aquatic and is considered challenging to monitor using conventional methods (Churchfield et al., 2000). Yet it is evident that eDNA represents a rapid and viable method for local detections (Yonezawa et al., 2020). Although the grey squirrel (*S. carolinensis*) is considered an arboreal species, it spends a significant proportion of its time foraging on the ground (more so than the red squirrel *S. vulgaris*) and is frequently detected here by eDNA (Figs 2 and 3).

As with other surveying techniques, behaviour and ecology can influence eDNA detection in natural water bodies (Harper et al., 2019; Williams et al., 2018). As previous studies have highlighted, carnivores are typically difficult to detect from water-based eDNA (Harper et al., 2019; Sales, McKenzie et al., 2020; Sales, Kaizer et al., 2020). It has been proposed that this is due to the fact that they are generally solitary, wide-ranging and may have less frequent contact with water bodies (e.g., animals might excrete/defecate on land more frequently) which could produce non-detections despite being present in the area/region (leading to potential false negatives from eDNA; Harper et al., 2019; Leempoel et al., 2020; Sales, McKenzie et al., 2020). Even when known or when regularly captured on camera traps in an area, eDNA has either infrequently, or completely failed to, detect species from this order (Harper et al., 2019; Sales, McKenzie et al., 2020). This study in particular has highlighted the intensity of sampling required to detect all the wild carnivores within a given area (Figs 2 and 4), even with semi-aquatic species such as the otter (*L. lutra*). Although all seven species within the order known in this geographic region were detected, three of these were only detected at a single site and no carnivore was detected on both the Colne and Blackwater (the red fox *V. vulpes* was additionally detected in the beaver experiment; Fig. S5; Table S2). Only the stoat (*M. erminea*) and American mink (*N. vison*) had comparatively higher detection probabilities (Fig. 3; Table S4).

This finding in relation to the American mink is note-worthy because the current study area is an active mink eradication zone due to the critical impacts this invasive species has had on local water vole populations. Browett et al. (2020) discussed the potential application of eDNA-based monitoring for the early detection of invasive mammals and given this mink eDNA detection on the periphery of the eradication zone, continuous monitoring using eDNA-based methods could be warranted to aid in keeping this region and others mink-free. The key for both early detection of invasive species and monitoring critically endangered species is the ability to detect species at low abundance (i.e. when a small number of individuals first colonize/invade an area). The number of individuals present in a system is clearly an important factor for eDNA detection (Williams et al., 2018). This may explain why some species listed as vulnerable such as the hedgehog *Erinaceus europaeus* (a species which has been declining in the UK; Mathews & Harrower, 2020) and locally rare hazel dormouse *Muscardinus avellanarius* (Dobson & Tansley, 2014) were not detected by eDNA (Table S2) and may require more targeted eDNA-based approaches (Priestley et al., 2021). The results from the beaver experiment may be particularly informative in this case however for early detection/detections at low abundance using eDNA. It is of course a semi-aquatic species but only two individuals were present in the enclosure and yet the species was still readily detected in multiple water replicates 300-400m downstream of the enclosure (Figs 1B and S5). It is clear that further experiments are required for a more complete understanding of the effects of species abundance along with transport and persistence of eDNA in relation to terrestrial and semi-aquatic mammals. This certainly presents unique challenges in comparison to studies involving species which are fully aquatic (Sales, McKenzie et al., 2020).

In addition to individual species detections, eDNA has the potential to improve our understanding of the dynamics of biodiversity in both time and space (Deiner et al., 2017). This is essential for the effective management of species and their ecosystems. The investigation of species richness and β-diversity patterns obtained from eDNA samples might provide insights into the underlying processes which structure communities (Deiner et al., 2017). Species assemblages can differ in two potential ways: the first, namely species turnover, which is based on the replacement of species between sites (i.e. substitution of one species in one site by a different one in the other site); and the second way, known as nestedness, refers to a pattern where there is species loss or gain, implying that different sites are strict subsets of richer ones (Baselga et al., 2012). In this study, dissimilarity (β-diversity) within both rivers was mainly driven by species turnover. This pattern was more evident in the Colne than Blackwater. This pattern indicates that the landscape surrounding each river presents a fairly distinct subset of species due to the high occurrence of species replacement. Contrasting to the pattern found within each river, a comparison between both river systems revealed a pattern consistent with higher nestedness-resultant dissimilarity. Nestedness occurs when assemblages of sites with fewer species tend to be a subset of the biotas from richer sites (Wright and Reeves, 1992). This result corroborates with the difference in species richness and composition between rivers (Fig. 5) and indicates that the Blackwater may represent a subset of the Colne in terms of the mammalian community surrounding it.

Different processes might be responsible for shaping community structure (Podani and Schmera, 2011, Gutiérrez-Cánovas et al., 2013). Partitioning β-diversity into its spatial turnover and nestedness components contributes to disentangle the processes underlying beta diversity and understand the putative drivers of community change (Baselga et al., 2010, Legendre and De Cáceres, 2013). For conservation purposes, this distinction is paramount for considering that both are antithetic processes that require distinct conservation strategies (Wright and Reeves, 1992). In a nestedness scenario, the prioritization of a small number of the richest sites could be considered, whereas for a species turnover scenario, a higher conservation effort of a large number of sites would be advised (Wright and Reeves, 1992). This assumption can be extrapolated to an eDNA sampling strategy for monitoring mammals from water taken from riverine systems. Here, by comparing both rivers for the purposes of monitoring mammals at a landscape-level, we demonstrate that the choice of sampling a single system in a nestedness context could lead to two different outcomes: sampling the richest system and obtaining the detection of most of the mammal diversity found in the area (e.g., sampling only the River Colne), or sampling the less diverse system and obtaining an underestimation due to the sampling effort directed towards detecting solely a subset of the biota from a richer river (e.g., sampling only the River Blackwater). Therefore, choosing to only sample a single river could have consequences for the inferred surrounding mammalian community. Just as importantly, the high turnover contribution within each river course might indicate that there is a low contribution of eDNA transport. eDNA longitudinal transport and diffusion would in theory lead to a higher nestedness within sites located upstream, leading to them being subsets from sites located further downstream in the river (because of eDNA accumulation). Preferentially sampling sites downstream (which may represent ‘eDNA reservoirs’; Sales et al., 2021) does not seem to represent an optimal sampling strategy for species which are not fully aquatic based on the results presented here.

This study demonstrates that eDNA metabarcoding from river-derived water is an efficient method for mapping mammalian distributions and diversity, and is highly capable of identifying the vast majority of the expected terrestrial and semi-aquatic species from a variety of mammalian orders within a short but intensive sampling period. We quantitatively demonstrate the effort required to capture different species within different orders and that considerations around individual species’ ecologies are important for eDNA-based monitoring of mammalian communities. This study provides the scope for using eDNA from rivers to quantify genuine differences in mammalian communities in their vicinity, and allows for the future incorporation of biotic and abiotic variables to understand the underlying factors behind these differences (Mariani et al., 2021). Adopting eDNA-based approaches for mammals would provide a reliable complement to ongoing national surveying efforts and one that could be readily adapted to involve citizen scientists.

## Acknowledgements

We would like to thank the Essex Wildlife Trust, Natural England and University of Salford for funding this study. Thanks to the landowners and river wardens who facilitated access to the sampling sites. We are grateful to Delphine Pouget for initial discussions about the applications of eDNA to monitor UK mammals. We thank Ilaria Coscia and Jake Jackman for sharing R scripts and providing advice on analyses.

## Author contributions

ADM, DT and NGS conceived and designed the study. HAB, EKB, JCP, NS, DT and ADM performed the sampling. HAB, EKB, JCP, SSB and NGS performed the laboratory work and HAB, LMG, NGS, JVL, PB and SSB the bioinformatics. LMG, HAB, NGS and ADM analysed the data. HAB, LMG, NGS and ADM wrote the manuscript, with all authors contributing to discussions and editing.

## Data Availability Statement

All bioinformatic steps and scripts can be found on github (https://github.com/McDevitt-Lab). Raw sequence data will be made publicly available upon peer-reviewed publication.

