## Supplementary Material for "Mapping differences in mammalian distributions and diversity using environmental DNA from rivers"

| **TABLE OF CONTENTS** | Page |
| --- | --- |
| **Table S1.**  Number of reads retained after each filtering step (after bioinformatics) to account for tag-switching, removing non-target MOTUs and MOTUs/reads originating from putative sequencing errors or contamination. | 3 |
| **Table S2.** Species (and the order to which they belong) detected in Essex, UK, including the records available in Matthews et al (2018) and species detected using eDNA metabarcoding for the river Colne, river Blackwater and for the beaver experiment. The number of sites of positive detection, IUCN status (England; IUCN, 2020) and whether they are naturalised (Nat) or non-native (NN; Crowley et al., 2020) are also specified. In bold, species that have been recorded in the sampling location but have not been detected by eDNA metabarcoding, and in red are species detected by eDNA but not included in the species list. | 4 |
| **Table S3.** Results obtained from Chao II estimator for the number of species detected by environmental DNA (eDNA) on the Rivers Colne and Blackwater combined (Total) and by each individual river separately. The 95% confidence intervals around these estimates are indicated (95% CI). | 5 |
| **Table S4**. Estimated site occupancies and detection probabilities for each species (with associated 95% confidence intervals in brackets) obtained from eDNA metabarcoding data (based on presence-absence data from five water replicates) from the River Colne and Blackwater separately and combined. | 6 |
| **Figure S1.** Environmental DNA sample site locations along the rivers Blackwater (B6–B15; A) and Colne (C1–C15; A), inside the beaver enclosure (B1–B3; B), downstream of the beaver enclosure (B4; B) and the large pond inlet (B5; B). | 8 |
| **Figure S2.** Examples of three environmental DNA (eDNA) sampling areas on the river Colne (A), river Blackwater (B) and inside beaver enclosure (C). Sampling methodology for eDNA is indicated in (C), where water was collected from the edge of the water body. | 9 |
| **Figure S3**. A bubble graph representing presence-absence and categorical values of proportional reads retained after bioinformatic analysis and conservative filtering for each wild mammal identified in each site on the River Colne (C1–C15). | 10 |
| **Figure S4.** A bubble graph representing presence-absence and categorical values of proportional reads retained after bioinformatic analysis and conservative filtering for each wild mammal identified in each site on the River Blackwater (B6–B15). | 11 |
| **Figure S5.** A bubble graph representing presence-absence and categorical values of proportional reads retained after bioinformatic analysis and conservative filtering for each wild mammal identified in each site in the beaver enclosure (B1–B3), downstream of the beaver enclosure (B4) and the large pond inlet (B5). | 12 |

**Table S1.**  Number of reads retained after each filtering step (after bioinformatics) to account for tag-switching, removing non-target MOTUs and MOTUs/reads originating from putative sequencing errors or contamination.

|  | **Library 1** | **Library 2** |
| --- | --- | --- |
| Total reads before filtering criteria | 5,500,594 | 6,520,512 |
| After accounting for tag-switching | 5,069,328 | 6,071,712 |
| After removing positive control | 5,069,256 | 6,071,609 |
| After removing negative controls | 4,906,193 | 5,665,106 |
| After removing all non-mammal reads | 3,888,233 | 4,223,874 |
| After removing human reads | 1,958,199 | 3,027,859 |
| After removing domestic animals (*Sus, Bos, Felis, Ovis* and *Canis*) | 614,467 | 1,923,338 |
| MOTU’s with minimum identity >0.98 and MOTUs with less than 5 reads removed | 588,776 | 1,858,913 |
| Tag-switching proportion | 0.00025 | 0.0004 |

**Table S2.** Species (and the order to which they belong) detected in Essex, UK, including the records available in Mathews et al (2018) and species detected using eDNA metabarcoding for the River Colne, River Blackwater and for the beaver experiment. The number of sites of positive detection, IUCN status (England; IUCN, 2020) and whether they are naturalised (Nat) or non-native (NN; Crawley et al., 2020) are also specified. In bold, species that have been recorded in the sampling location but have not been detected by eDNA metabarcoding, and in red are species detected by eDNA but not included in the species list.

| **Order** | **Scientific name** | **Common name** | **Order** | **IUCN status (England) *** | **Colne** | **Blackwater** | **Beaver** |
| --- | --- | --- | --- | --- | --- | --- | --- |
| *Carnivora* | *Neovison vison* | American mink | Carnivora | NN | 1 | 0 | 0 |
|  | *Lutra lutra* | European otter | Carnivora | LC | 2 | 0 | 0 |
|  | *Mustela nivalis* | Least weasel | Carnivora | LC | 1 | 0 | 0 |
|  | *Mustela erminea* | Stoat | Carnivora | LC | 2 | 0 | 0 |
|  | *Mustela putorius* | European polecat | Carnivora | LC | 0 | 1 | 0 |
|  | *Meles meles* | European badger | Carnivora | LC | 7 | 0 | 0 |
|  | *Vulpes vulpes* | Red fox | Carnivora | LC | 1 | 0 | 1 |
| Rodentia | *Arvicola amphibius* | European water vole | Rodentia | EN | 9 | 4 | 2 |
|  | *Myodes glareolus* | Bank vole | Rodentia | LC | 5 | 3 | 4 |
|  | *Microtus agrestis* | Field vole | Rodentia | LC | 13 | 8 | 3 |
|  | *Apodemus sylvaticus* | Wood mouse | Rodentia | LC | 1 | 5 | 3 |
|  | *Apodemus flavicollis* | Yellow-necked mouse | Rodentia | LC | 1 | 1 | 0 |
|  | *Rattus norvegicus* | Brown rat | Rodentia | NN | 13 | 5 | 2 |
|  | *Sciurus carolinensis* | Grey squirrel | Rodentia | NN | 5 | 7 | 3 |
|  | *Sciurus vulgaris* | Red squirrel | Rodentia | EN | 1 | 0 | 0 |
|  | *Castor fiber* | Eurasian beaver | Rodentia | CE | 0 | 0 | 4 |
|  | ***Muscardinus avellanarius*** | **Hazel dormouse** | Rodentia | VU | 0 | 0 | 0 |
|  | ***Micromys minutus*** | **Harvest mouse** | Rodentia | LC | 0 | 0 | 0 |
|  | ***Mus musculus*** | **House mouse** | Rodentia | Nat | 0 | 0 | 0 |
| Artiodactyla | *Muntiacus reevesi* | Reeves's muntjac | Artiodactyla | NN | 2 | 5 | 4 |
|  | *Dama dama* | Fallow deer | Artiodactyla | Nat | 5 | 0 | 0 |
|  | *Capreolus capreolus* | Roe deer | Artiodactyla | LC | 1 | 0 | 0 |
|  | ***Cervus elaphus*** | **Red deer** | Artiodactyla | LC | 0 | 0 | 0 |
| Eulipotyphla | *Neomys fodiens* | Water shrew | Eulipotyphla | LC | 5 | 2 | 1 |
|  | *Sorex minutus* | Pygmy shrew | Eulipotyphla | LC | 2 | 0 | 1 |
|  | *Sorex araneus* | Common shrew | Eulipotyphla | LC | 4 | 0 | 0 |
|  | *Talpa europaea* | European mole | Eulipotyphla | LC | 1 | 3 | 0 |
|  | ***Erinaceus europaeus*** | **Hedgehog** | Eulipotyphla | VU | 0 | 0 | 0 |
| Lagomorpha | *Lepus europaeus* | Brown hare | Lagomorpha | Nat | 4 | 0 | 0 |
|  | *Oryctolagus cuniculus* | European rabbit | Lagomorpha | Nat | 7 | 3 | 1 |

*CE - critically endangered, EN - endangered, VU - vulnerable, LC - least concern, Nat - naturalised, NN - non-native

**Table S3.** Results obtained from Chao II estimator for the number of species detected by environmental DNA (eDNA) on the Rivers Colne and Blackwater combined (Total) and by each individual river separately. The 95% confidence intervals around these estimates are indicated (95% CI).

| **Type** | **Species Richness** | **Chao II** | **95% CI** |
| --- | --- | --- | --- |
| Total | 24 | 26.88 | 24.48-41.43 |
| Colne | 23 | 28.23 | 24.07-48.54 |
| Blackwater | 12 | 12.45 | 12.03-19.58 |

**Table S4**. Estimated site occupancies and detection probabilities for each species (with associated 95% confidence intervals in brackets) obtained from eDNA metabarcoding data (based on presence-absence data from five water replicates) from the River Colne and Blackwater separately and combined.

| **Species** | **Order** | **Occupancy** | | | **Detection Probability** | | |
| --- | --- | --- | --- | --- | --- | --- | --- |
|  |  | Combined | Colne | Blackwater | Combined | Colne | Blackwater |
| *Neovison vison* | Carnivora | 0.04 (0.01-0.24) | 0.07 (0.01-0.35) | - | 0.80 (0.31-0.97) | 0.80 (0.31-0.97) | - |
| *Lutra lutra* | Carnivora | 0.12 (0.02-0.46) | 0.20 (0.03-0.65) | - | 0.20 (0.03-0.66) | 0.20 (0.03-0.66) | - |
| *Mustela nivalis* | Carnivora | 0.98 (0-1.00) | 0.99 (0-1.00) | - | 0.01 (0-0.11) | 0.01 (0-0.12) | - |
| *Mustela erminea* | Carnivora | 0.08 (0.02-0.28) | 0.14 (0.03-0.42) | - | 0.48 (0.19-0.78) | 0.48 (0.19-0.78) | - |
| *Mustela putorius* | Carnivora | 0.98 (0-1.00) | - | 0.99 (0-1.00) | 0.01 (0-0.11) | - | 0.02 (0-0.15) |
| *Meles meles* | Carnivora | 0.38 (0.15-0.67) | 0.63 (0.21-0.91) | - | 0.24 (0.10-0.46) | 0.24 (0.10-0.46) | - |
| *Vulpes vulpes* | Carnivora | 0.98 (0-1.00) | 0.99 (0-1.00) | - | 0.01 (0-0.11) | 0.01 (0-0.12) | - |
| *Arvicola amphibius* | Rodentia | 0.65 (0.34-0.87) | 0.76 (0.28-0.96) | 0.49 (0.15-0.84) | 0.27 (0.16-0.43) | 0.27 (0.14-0.46) | 0.28 (0.11-0.57) |
| *Myodes glareolus* | Rodentia | 1.00 (0-1.00) | 0.9 (0-1.00) | 1.00 (0-1.00) | 0.07 (0.04-0.14) | 0.09 (0.01-0.42) | 0.06 (0.02-0.17) |
| *Microtus agrestis* | Rodentia | 0.94 (0.36-1.00) | 0.90 (0.54-0.99) | 1.00 (0-1.00) | 0.36 (0.26-0.47) | 0.48 (0.35-0.61) | 0.20 (0.11-0.33) |
| *Apodemus sylvaticus* | Rodentia | 0.31 (0.12-0.59) | 0.99 (0-1.00) | 0.60 (0.21-0.90) | 0.26 (0.11-0.49) | 0.01 (0-0.12) | 0.30 (0.13-0.55) |
| *Apodemus flavicollis* | Rodentia | 1.00 (0-1.00) | 0.99 (0-1.00) | 0.99 (0-1.00) | 0.02 (0-0.07) | 0.01 (0-0.12) | 0.02 (0-0.15) |
| *Rattus norvegicus* | Rodentia | 0.75 (0.53-0.89) | 0.91 (0.51-0.99) | 0.51 (0.23-0.79) | 0.49 (0.38-0.60) | 0.46 (0.33-0.60) | 0.55 (0.35-0.74) |
| *Sciurus carolinensis* | Rodentia | 0.63 (0.31-0.86) | 0.9 (0-1.00) | 0.79 (0.31-0.97) | 0.26 (0.14-0.42) | 0.09 (0.01-0.42) | 0.36 (0.20-0.55) |
| *Sciurus vulgaris* | Rodentia | 0.98 (0-1.00) | 0.99 (0-1.00) | - | 0.01 (0-0.11) | 0.01 (0-0.12) | - |
| *Muntiacus reevesi* | Artiodactyla | 0.29 (0.15-0.50) | 1.00 (0-1.00) | 0.51 (0.23-0.78) | 0.46 (0.29-0.64) | 0.03 (0.01-0.10) | 0.59 (0.39-0.77) |
| *Dama dama* | Artiodactyla | 0.24 (0.09-0.49) | 0.41 (0.15-0.72) | - | 0.30 (0.13-0.55) | 0.30 (0.13-0.54) | - |
| *Capreolus capreolus* | Artiodactyla | 0.98 (0-1.00) | 0.99 (0-1.00) | - | 0.01 (0-0.11) | 0.01 (0-0.12) | - |
| *Neomys fodiens* | Eulipotyphla | 0.39 (0.15-0.69) | 0.46 (0.14-0.81) | 0.29 (0.04-0.79) | 0.23 (0.10-0.45) | 0.24 (0.09-0.50) | 0.20 (0.03-0.66) |
| *Sorex minutus* | Eulipotyphla | 0.12 (0.02-0.46) | 0.20 (0.03-0.65) | - | 0.20 (0.03-0.66) | 0.20 (0.03-0.66) | - |
| *Sorex araneus* | Eulipotyphla | 0.24 (0.07-0.58) | 0.40 (0.10-0.80) | - | 0.20 (0.06-0.51) | 0.20 (0.06-0.51) | - |
| *Talpa europaera* | Eulipotyphla | 0.20 (0.07-0.46) | 0.99 (0-1.00) | 0.34 (0.10-0.69) | 0.28 (0.11-0.57) | 0.01 (0-0.12) | 0.36 (0.14-0.65) |
| *Lepus europaeus* | Lagomorpha | 0.24 (0.07-0.58) | 0.40 (0.10-0.80) | - | 0.20 (0.06-0.51) | 0.20 (0.06-0.51) | - |
| *Oryctolagus cuniculus* | Lagomorpha | 0.45 (0.25-0.67) | 0.54 (0.25-0.81) | 0.32 (0.10-0.65) | 0.37 (0.23-0.53) | 0.33 (0.17-0.54) | 0.44 (0.20-0.71) |

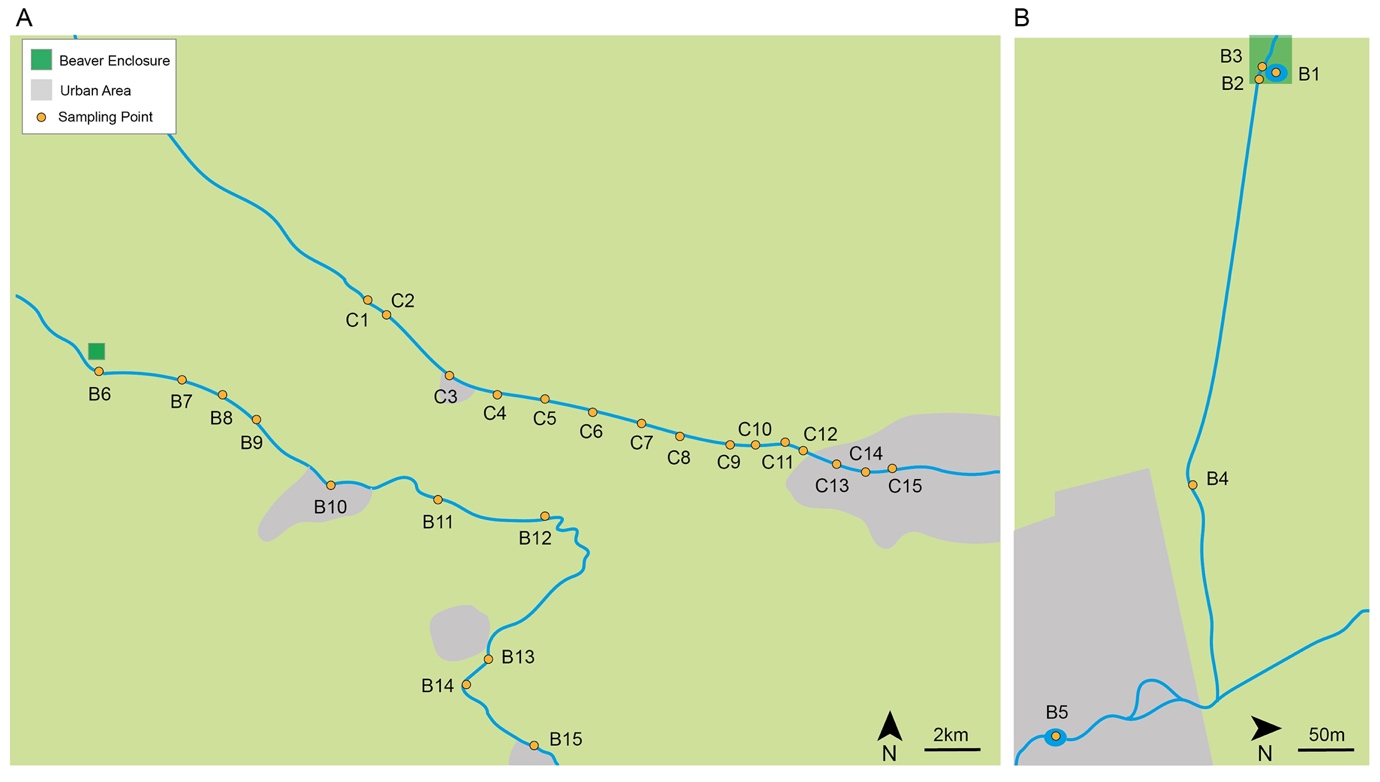

**Figure S1.** Environmental DNA sampling locations along the rivers Blackwater (B6–B15; A) and Colne (C1–C15; A), inside the beaver enclosure (B1–B3; B), downstream of the beaver enclosure (B4; B) and the large pond inlet (B5; B).

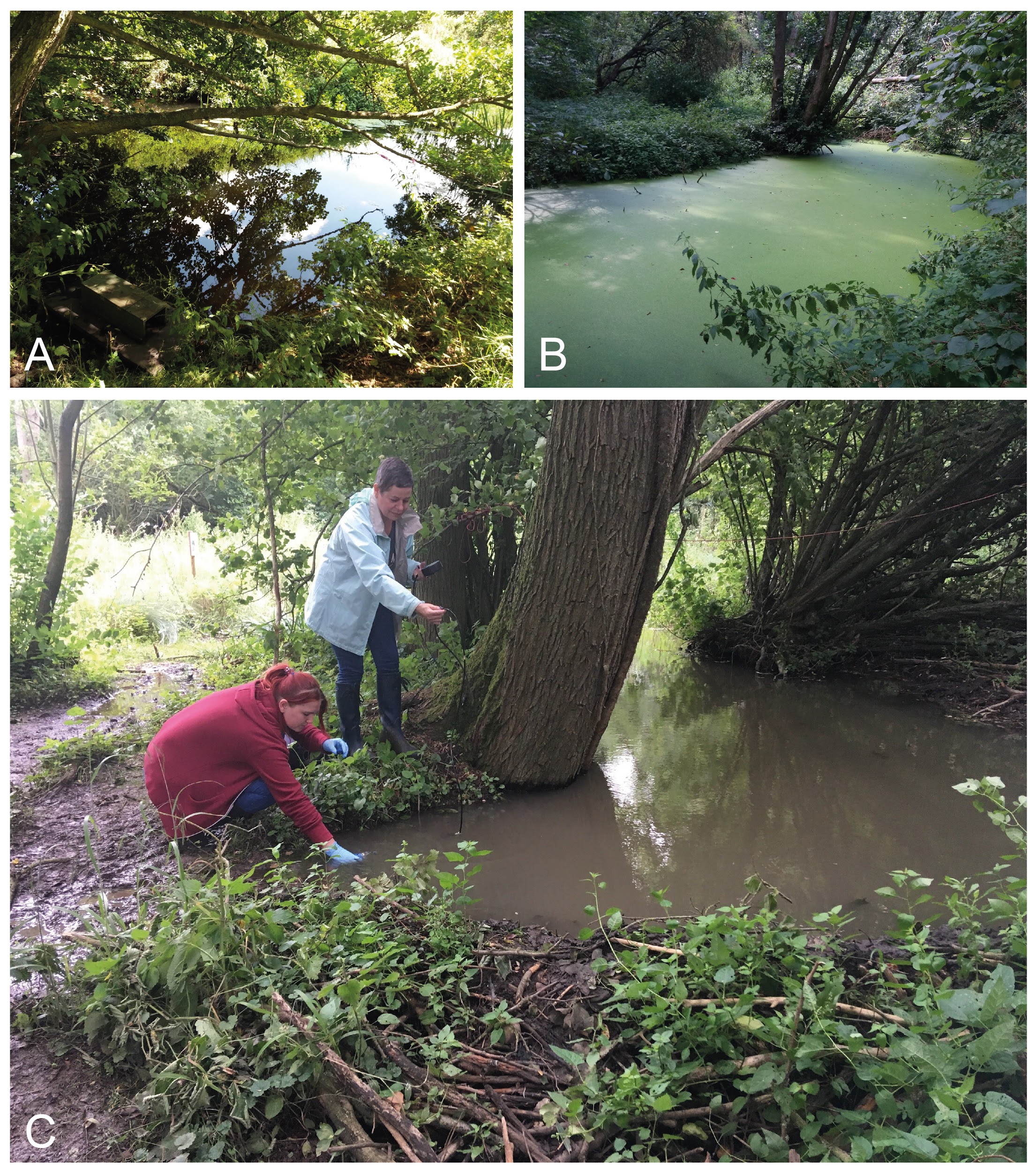

**Figure S2.** Examples of three environmental DNA (eDNA) sampling areas on the river Colne (A), river Blackwater (B) and inside the beaver enclosure (C). Sampling methodology for eDNA is indicated (C), where water was collected from the edge of the water body.

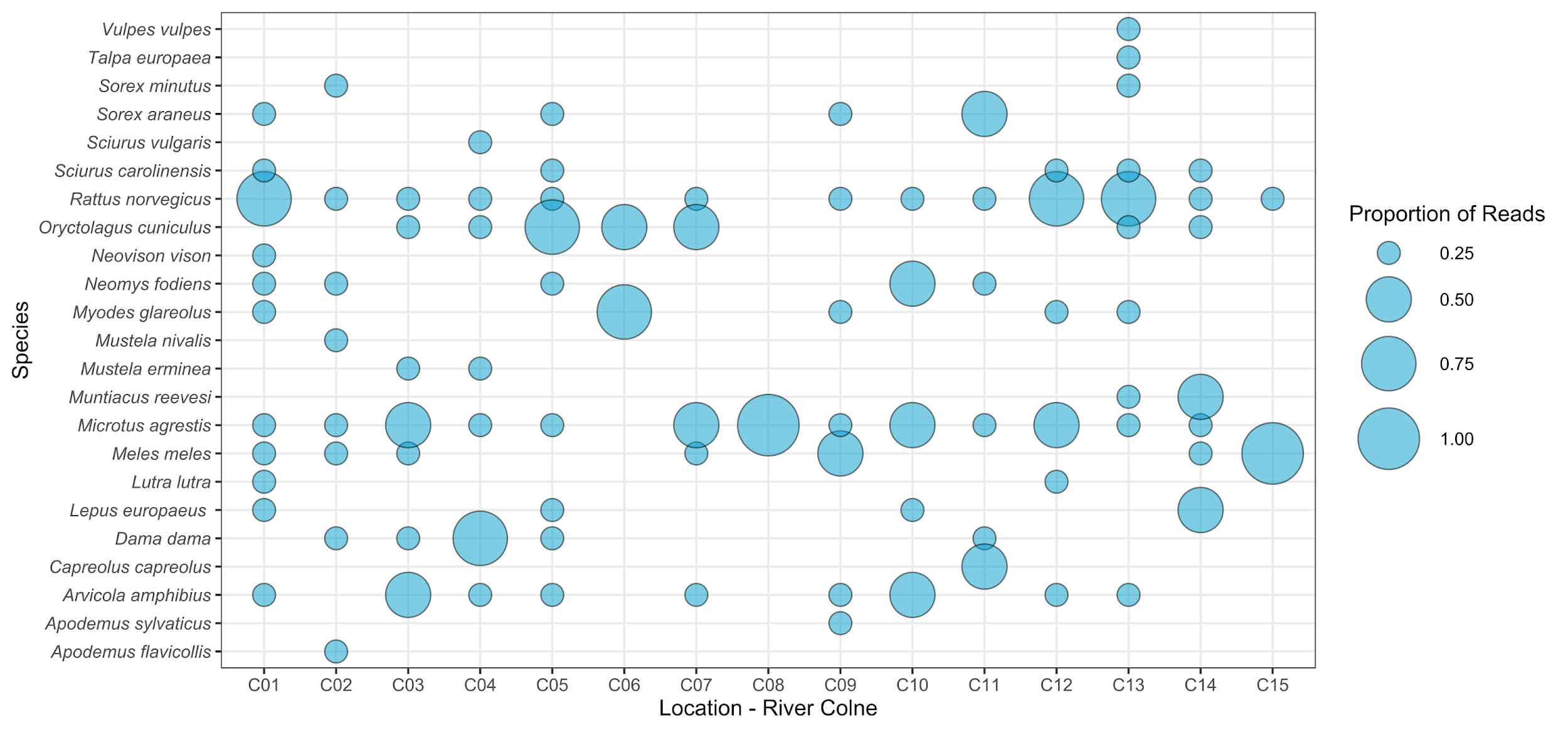

**Figure S3**. A bubble graph representing presence-absence and categorical values of proportional read counts retained after bioinformatic analysis and filtering for each wild mammal identified in each site on the River Colne (C1–C15).

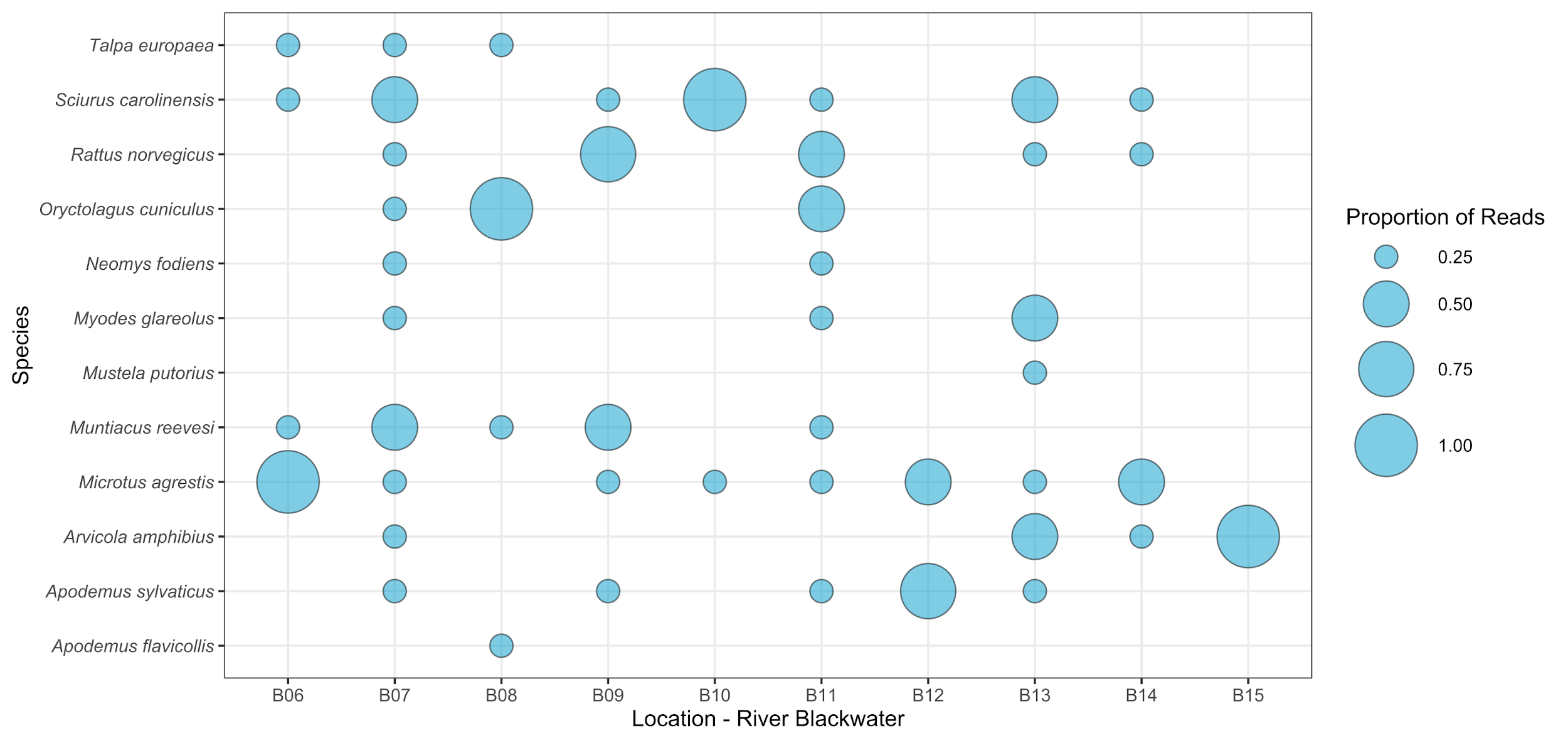

**Figure S4.** A bubble graph representing presence-absence and categorical values of proportional read counts retained after bioinformatic analysis and filtering for each wild mammal identified in each site on the River Blackwater (B6–B15).

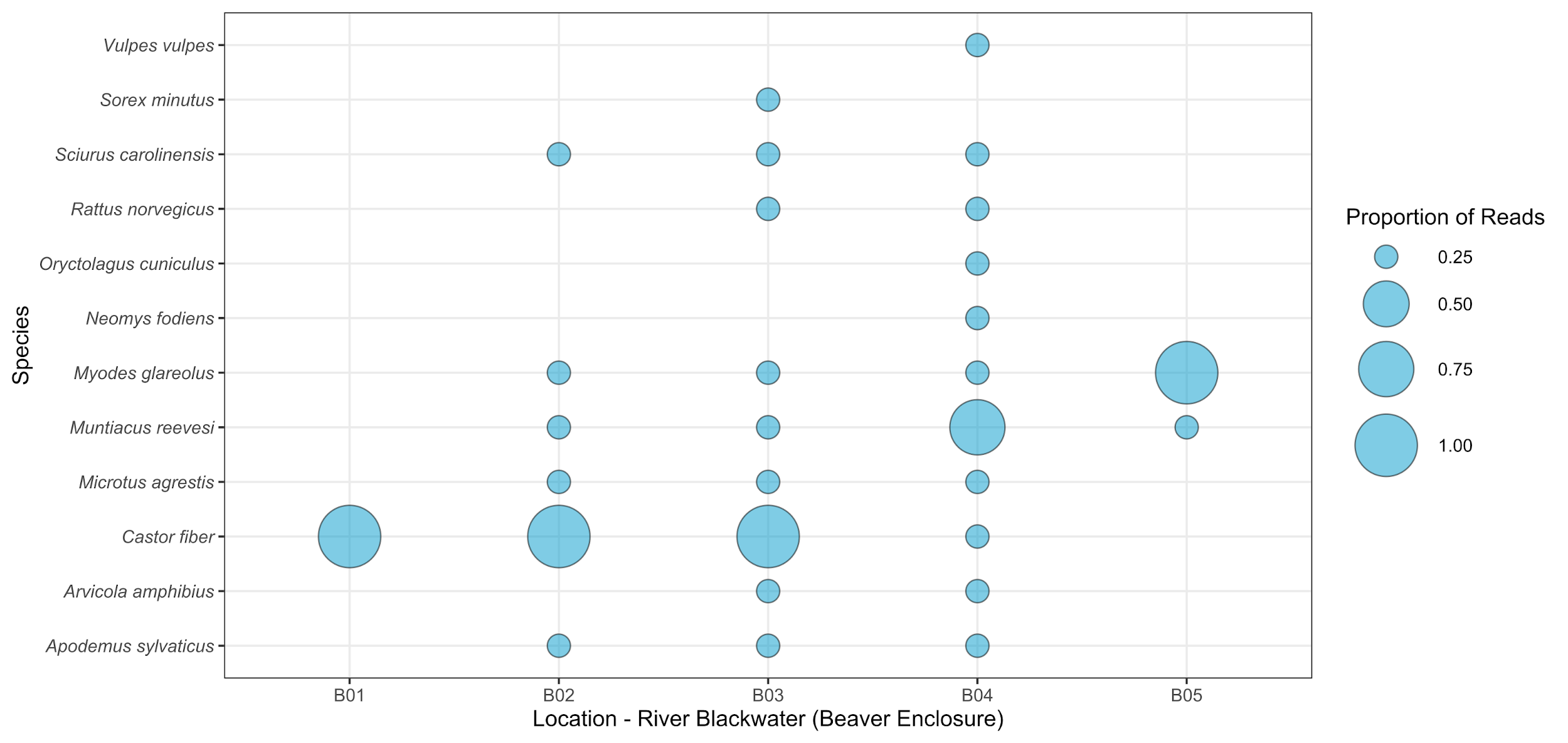

**Figure S5.** A bubble graph representing presence-absence and categorical values of proportional read counts retained after bioinformatic analysis and filtering for each wild mammal identified in each site in the beaver enclosure (B1–B3), downstream of the beaver enclosure (B4) and the large pond inlet (B5).
